# Simplitigs as an efficient and scalable representation of de Bruijn graphs

**DOI:** 10.1101/2020.01.12.903443

**Authors:** Karel Břinda, Michael Baym, Gregory Kucherov

**Author notes:** These authors contributed equally to this work (Michael Baym, Gregory Kucherov).

## Abstract

De Bruijn graphs play an essential role in computational biology. However, despite their widespread use, they lack a universal scalable representation suitable for different types of genomic data sets. Here, we introduce simplitigs as a compact, efficient and scalable representation and present a fast algorithm for their computation. On examples of several model organisms and two bacterial pan-genomes, we show that, compared to the best existing representation, simplitigs provide a substantial improvement in the cumulative sequence length and their number, especially for graphs with many branching nodes. We demonstrate that this improvement is amplified with more data available. Combined with the commonly used Burrows-Wheeler Transform index of genomic sequences, simplitigs substantially reduce both memory and index loading and query times, as illustrated with large-scale examples of GenBank bacterial pan-genomes.

## Background

Advances in DNA sequencing started the golden age of biology when previously unobservable phenomena can be studied on an unprecedented scale. However, sequencing capacity has been growing faster than computer performance and memory, and also faster than available human resources. Today large amounts of sequence data are available. Consequently, traditional sequence-based representations and sequence alignment-based techniques [1–3] have become less suitable for real-life scenarios due to the space- and time-complexities they impose and their inefficiency in handling polymorphisms.

De Bruijn graphs provide an elegant solution for genomic data representation. They build on top of the concept of *k*-mers, which are substrings of a fixed length *k* of the genomic strings to be represented, such as sequencing reads, genomes, or transcriptomes. For a given *k*-mer set, the corresponding de Bruijn graph is a directed graph with the *k*-mers being vertices and *k* − 1 long overlaps between pairs of these *k*-mers indicating edges (Methods). There is an obvious correspondence between *k*-mer sets and de Bruijn graphs, and we can use both terms interchangeably. If *k* is chosen appropriately, de Bruijn graphs capture substantial information about the sequenced molecules as these correspond to some walks in the graph.

The use of de Bruijn graphs is ubiquitous in sequence analysis. Genome assembly uses the property that sequenced molecules form paths [4–6], which is exploited in numerous modern assemblers [7–12]. On the other hand, alignment-free sequence comparison follows the idea that similar sequences share common *k*-mers, and comparing de Bruijn graphs provides thus a good measure of sequence similarity [13]. This involves applications of de Bruijn graphs to variant calling and genotyping [14–18], transcript abundance estimation [19], and metagenomic classification [20–23]. In the latter application, *k*-mer-based classifiers perform best among all classifiers in inferring abundance profiles [24], which suggests that de Bruijn graphs truthfully approximate the graph structures of bacterial pan-genomes, even if constructed from noisy assemblies from incomplete databases. Even if more advanced pan-genome graph representations are available, such as variation graphs [25], de Bruijn graphs with large *k*-mer lengths are still useful for indexing [26,27].

As de Bruijn graphs are one of the primary data structures in much of sequence analysis, the efficiency of many algorithms is directly tied to the efficiency of computation and representation of the graph. De Bruijn graphs can be readily computed through a scan of the datasets including the raw reads, genomes or multiple sequence files. In practice, such a scan often consists in *k*-mer counting as this allows efficient denoising of the graph, e.g., by removing low-frequency *k*-mers corresponding to sequencing errors in the reads. Algorithms for *k*-mer counting have been extensively studied and many well-engineered software solutions are available [28–36].

On the other hand, de Bruijn graph representations have received much less attention. The most commonly used representation are unitigs, which are strings resulting from compaction of *k*-mers along maximal paths with non-branching nodes [37,38]. Unitigs have many advantages: the representation is “textual”, in the form of a set of sequences that contain each *k*-mer exactly once while preserving graph topology. However, unitigs impose large resource overhead for many types of de Bruijn graphs and do not scale well when a lot of variation is included. Specifically, with a high proportion of branching nodes, unitigs become fragmented, in an extreme case up to the level of individual *k*-mers. Subsequently, unitig computation and storage may require inappropriately large resources and become prohibitive in variation-heavy applications, e.g., in bacterial pan-genomics.

In this paper, we propose simplitigs as a compact, efficient and scalable representation of de Bruijn graphs. Simplitigs generalize the unitig representation by relaxing the restriction of stopping at branching nodes. We present an algorithm for rapid simplitig computation from a *k*-mer set and implement it in a tool called ProphAsm. ProphAsm proceeds by loading a *k*-mer set into memory and a greedy enumeration of maximal vertex-disjoint paths in the associated de Bruijn graph. We use ProphAsm to evaluate the improvement of simplitigs over unitigs, in terms of two main characteristics: the cumulative sequence length (CL) and the number of sequences (NS). We demonstrate that greedily computed simplitigs are close to theoretical bounds in practical applications and provide, compared to unitigs, a substantial improvement in memory requirements and speed in applications such as *k*-mer matching.

## Results

### The concept of simplitigs

We developed the concept of simplitigs to efficiently represent de Bruijn graphs of sequence data (**Fig. 1**). Simplitigs are a generalization of unitigs and correspond to spellings of vertex-disjoint paths covering a given de Bruijn graph (**Fig. 1a**, Methods). Consequently, maximal simplitigs are such simplitigs where no two simplitigs can be merged by a (*k*−1) overlap (Methods). Note that unitigs and *k*-mers are also simplitigs, but not maximal in general (**Fig. 1b**). The main conceptual difference between maximal simplitigs and maximal unitigs is that simplitigs are not limited by branching nodes, which allows for further compaction, with a benefit increasing proportionally to the amount of branching nodes in the graph.

**Fig. 1.**
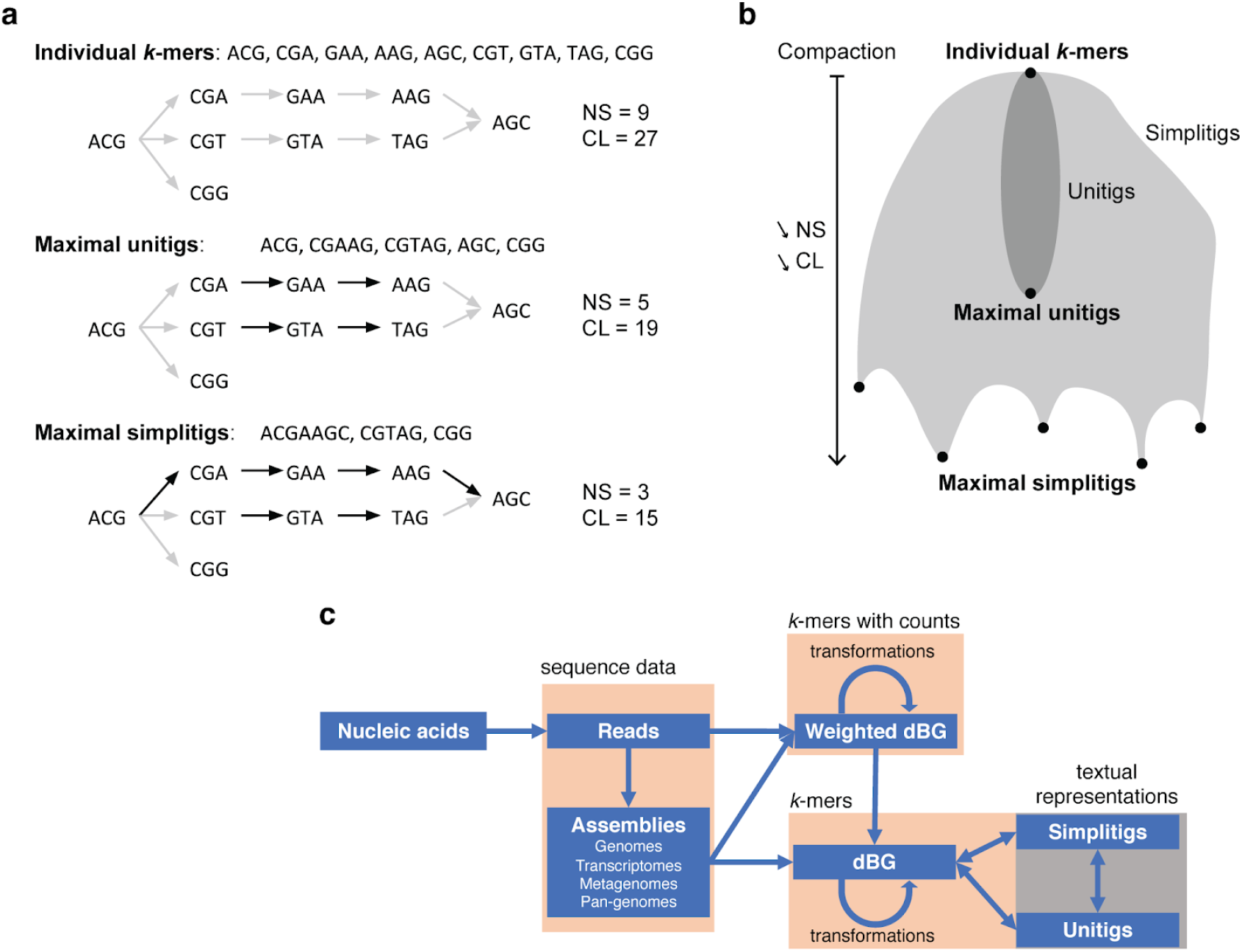
Overview of the simplitig approach. **a** Textual representations of *k*-mer sets ordered by the degree of compaction: individual *k*-mers, maximal unitigs, and maximal simplitigs. Every component of a simplitig subgraph (black arrows) of the de Bruijn graph (all arrows) corresponds to a path, and its spelling constitutes a simplitig (Methods). **b** Scheme of all possible simplitig representations according to the degree of compaction. While unitigs (dark gray area) correspond to compaction along non-branching nodes in the associated de Bruijn graph, simplitigs (gray area) can also contain branching nodes. Every step of compaction decreases the number of sequences (NS) and their cumulative length (CL) by 1 and by *k*−1, respectively. Maximal simplitigs may not be determined uniquely; the simplitig representation can have multiple local optima, depending on which edges were selected at the branching nodes. **c** The workflow of simplitigs. Simplitigs represent de Bruijn graphs and carry implicitly the same information as unitigs. De Bruijn graphs are usually computed from either assemblies or weighted de Bruijn graphs. Weighted de Bruijn graphs are typically obtained by *k*-mer counting and allow removing noise, e.g., low-frequency *k*-mers, which frequently originate in sequencing errors.

To compare simplitig and unitig representations, we created a benchmarking procedure based on the two characteristics: the number of sequences (NS) and their cumulative length (CL) (Methods, example in **Fig. 2**). While NS determines the number of records to be kept in memory, CL largely determines the total memory needed. NS and CL are readily bounded from below by one and by the number of *k*-mers, respectively, and they are also tightly connected ((eq 1) in Methods). As every step of compaction decreases both NS and CL (**Fig. 1b**, Methods), we can optimize them jointly. However, finding an optimal simplitig representation translates to the vertex disjoint path coverage problem. While NP-hard for general graphs (by reduction from the well-known NP-hard problem of computing a Hamiltonian path), the problem may be tractable for observed de Bruijn graphs (Methods).

**Fig. 2.**
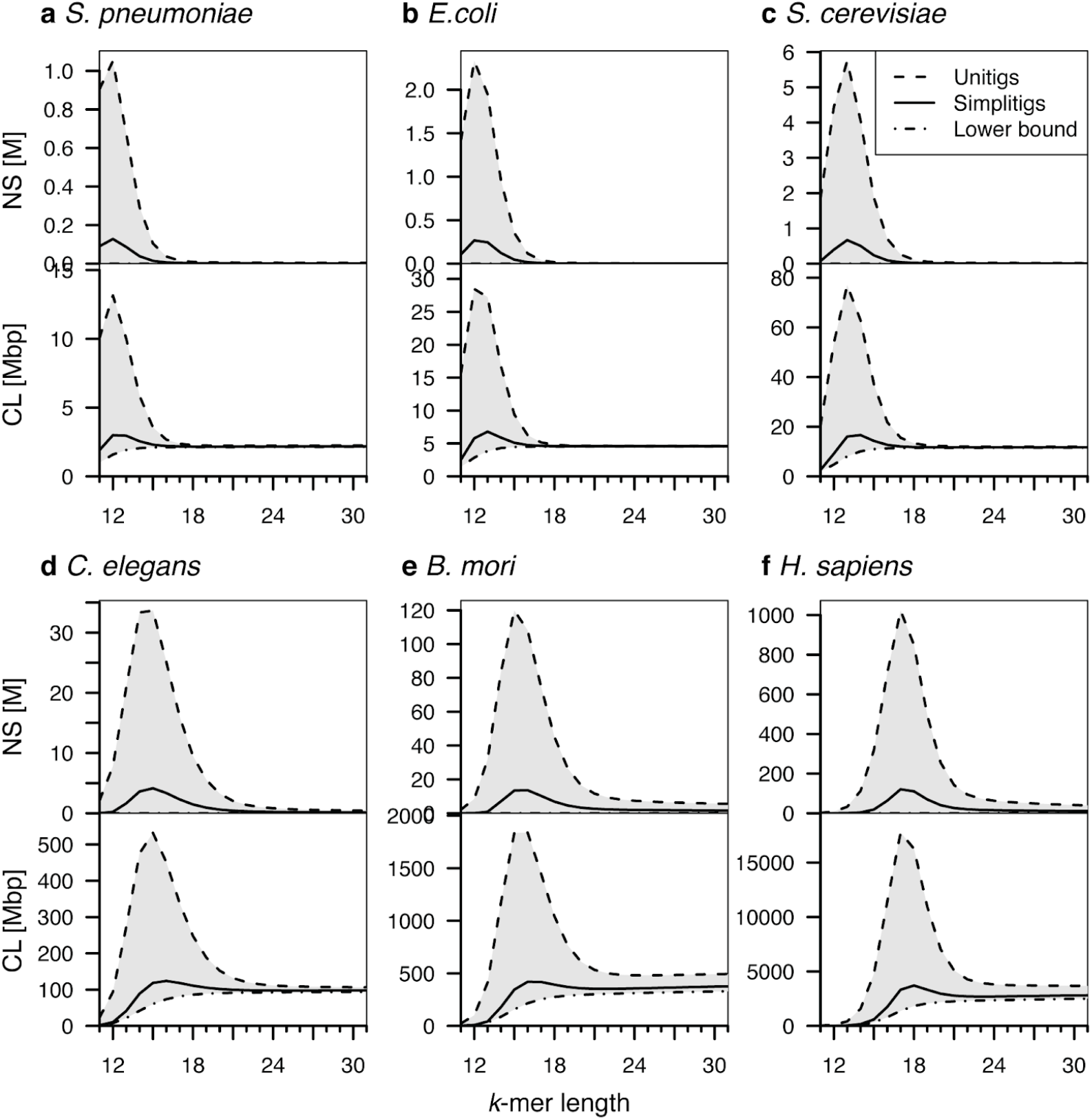
Comparison of the simplitig and unitig representations for selected model organisms and a range of *k*-mers. The number of sequences (NS, millions) and their cumulative length (CL, megabase pairs) for both representations of six model organisms ordered by their genome size: **a** *Streptococcus pneumoniae*, 2.22 Mbp; **b** *Escherichia coli*, Mbp; **c** *Saccharomyces cerevisiae*, 12.2 Mbp; **d** *Caenorhabditis elegans*, 100 Mbp; **e** *Bombyx mori*, 482 Mbp; and **f** *Homo sapiens*, 3.21 Gbp. The CL lower bound corresponds to the number of *k*mers. Full results are available in **Additional File 1**.

Since practical applications do not require optimal simplitigs, we prioritized speed and designed a greedy algorithm for their rapid computation (**Alg. 1**, Methods). In an iterative fashion, the algorithm selects an arbitrary *k*-mer as a seed of a new simplitig and keeps extending it forwards and then backwards as long as possible, while removing the already used *k*-mers from the set. This process is repeated until all *k*-mers are covered. Loading *k*-mers into memory and simplitig computation are linear in the length of the input and the number of *k*-mers, respectively, and the memory footprint is linear in the number of *k*-mers. We implemented **Alg. 1** in a program called ProphAsm, available at https://github.com/prophyle/prophasm.

**Alg. 1.**
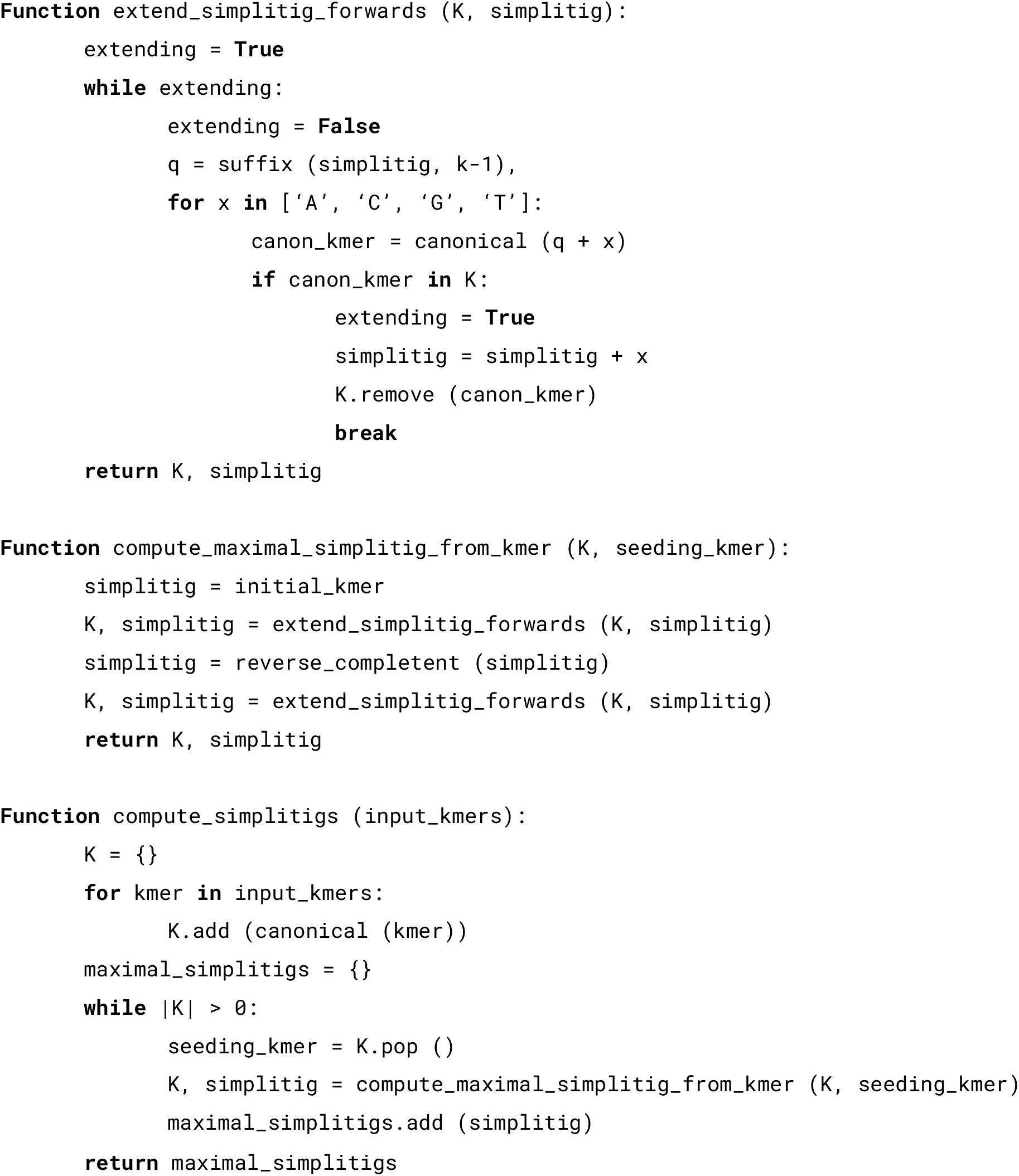
Greedy computation of maximal simplitigs for a *k*-mer set or a de Bruijn graph. In an iterative fashion, the algorithm draws a *k*-mer from the set of canonical *k*-mers *K*, uses it as a seed of a new simplitig, and then keeps extending the simplitig forwards as long as possible, and then backwards, while removing the already used canonical *k*-mers from *K*.

### Simplitigs of model organisms

We first evaluated simplitig and unitig representations on assemblies of six model organisms (**Fig. 2**). As different applications of de Bruijn graphs call for different *k*-mer lengths, we sought to characterize the NS and CL scaling for both representations with *k* growing, as well as the effect of the species’ genome size. Therefore, selected model organisms were evaluated in an increasing order of the genome size and benchmarked for both representations on a range of *k*-mer lengths corresponding to common alignment-free-based applications [19,20,39].

We observe that simplitigs always provide substantially better performance than unitigs (**Fig. 2**). In particular, they quickly approach the theoretical lower bounds for both characteristics tested. Every data set has a range of *k*-mer lengths where the difference between simplitigs and unitigs is very large, and after a certain threshold, the difference almost vanishes. While for short genomes this threshold is located at smaller *k*-mer lengths than those typically used in alignment-free applications (e.g., *k* ≈ 17 for *E. coli*), for bigger genomes this threshold has not been attained on the tested range and seems to be substantially shifted towards large *k*-mers (e.g., *B. mori*).

Interestingly, maxima of the NS and CL values for both representations occur very close to the value *k* = *log*_4_*G*, where *G* is the genome size (**Fig. 2**). This is readily explained by edge saturation: for values of *k* up to *log*_4_*G*, an overwhelming fraction of all 4^*k*^ *k*-mers belong to the genome, which makes the de Bruijn graph branch at nearly every node. As a consequence, unitigs are then essentially reduced to individual *k*-mers and their number grows exponentially whereas simplitigs stay compact on the whole range of *k*-mer lengths. Starting from *k* = *log*_4_*G* the graph starts to form longer non-branching paths, which drives down the NS and CL of unitigs, and they approach those of simplitigs. However, the difference between simplitigs and unitigs in their count and length may stay considerable even for larger values of *k*, especially in case of large eukaryotic genomes.

### Performance assessment

We then analyzed the computational resources that had been used to compute the simplitigs using ProphAsm and the unitigs using BCALM (**Fig. 3**). Both representations were computed in the environment of a computational cluster, with individual experiments deployed as parallel jobs using SLURM (Methods). Even though we primarily focused on the total CPU time, we also tested BCALM using 4 threads to evaluate the effect of parallelization.

**Fig. 3.**
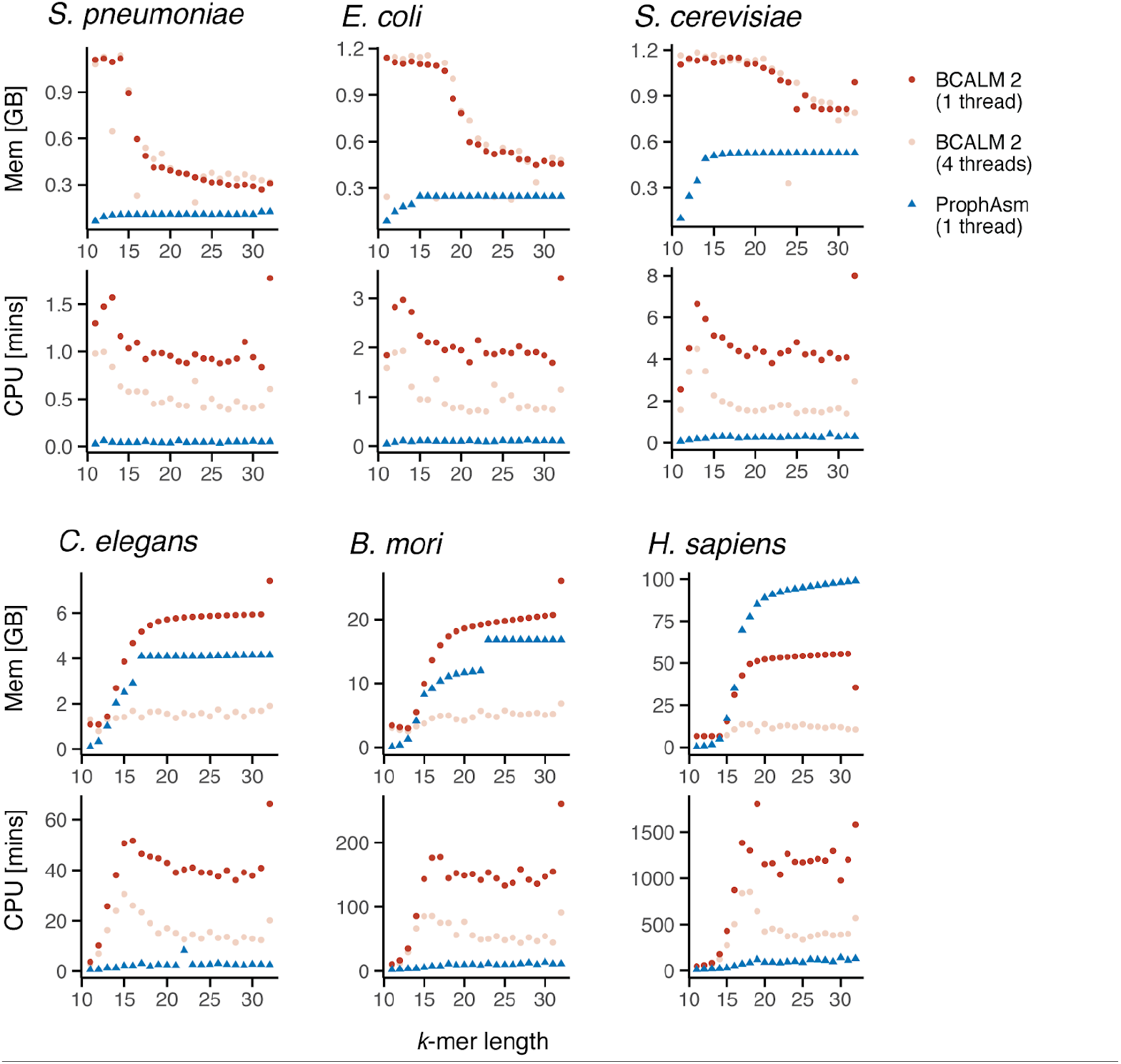
Comparison of CPU time and memory consumption of ProphAsm and BCALM. Resources to compute unitigs using BCALM (using one and four threads) and simplitigs using ProphAsm (using one thread) of the six model organisms. Full results are available in **Additional File 2**.

We observe that throughout our experiments ProphAsm outcompeted BCALM, both in terms of the memory consumption and the CPU time, even if BCALM was run with 4 threads (**Fig. 3**). The only exception was the memory consumption for *H. sapiens*, suggesting that BCALM may be more memory-efficient for large genomes. Importantly, ProphAsm used a consistent amount of resources: memory 38–51 bytes of RAM per *k*-mer in the dataset, a limited CPU time and no additional disk space. On the other hand, BCALM’s resource usage was less predictable and less consistent across experiments and required frequent trial-and-error resource adjustments and re-running (Methods).

Besides the comparatively high memory and CPU time requirements, the most challenging resource was the disk space. In order to fit within the available disk capacity, we reduced the BCALM consumption using the ‘-maxdisk’ parameter. However, despite that we requested BCALM to use no more than 30 GB per experiment, a random manual inspection revealed a substantially higher use – for instance, 116 GB of disk space for *H. sapiens* with *k* = 17 and 4 threads. The extensive disk space consumption also disabled a similar comparison on a desktop computer.

Overall, the better performance of ProphAsm can be explained by simplitigs being fundamentally easier to compute than unitigs and by BCALM being optimized for a particular class of de Bruijn graphs that emerge in assembly-like applications. As the ProphAsm resource usage is highly predictable, it appears to be more suitable for the use on clusters in many parallel instances, unless the number of *k*-mers in the dataset exceeds a critical threshold determined by the available RAM.

### Simplitigs of bacterial pan-genomes

We then sought to evaluate the impact of additional variation in a de Bruijn graph (**Fig. 4**). Such variation may originate in polymorphisms, varying gene content in a population of genomes that are represented jointly, in haplotypes of viral quasispecies, or in sequencing errors in case of graphs constructed directly from sequencing reads. In all these cases, many nodes of the de Bruijn graph become branching and new paths emerge. To model gradually increasing variation, we used bacterial pan-genomes with different levels of sampling. Given the high diversity and variability of bacteria, de Bruijn graphs provide a convenient option for computational pan-genomes [40]. Such pan-genomes can be constructed from draft assemblies or even directly from sequencing reads, and thanks to bacterial genomes being short and haploid, the information captured by the graphs is sufficient for many analyses.

**Fig. 4.**
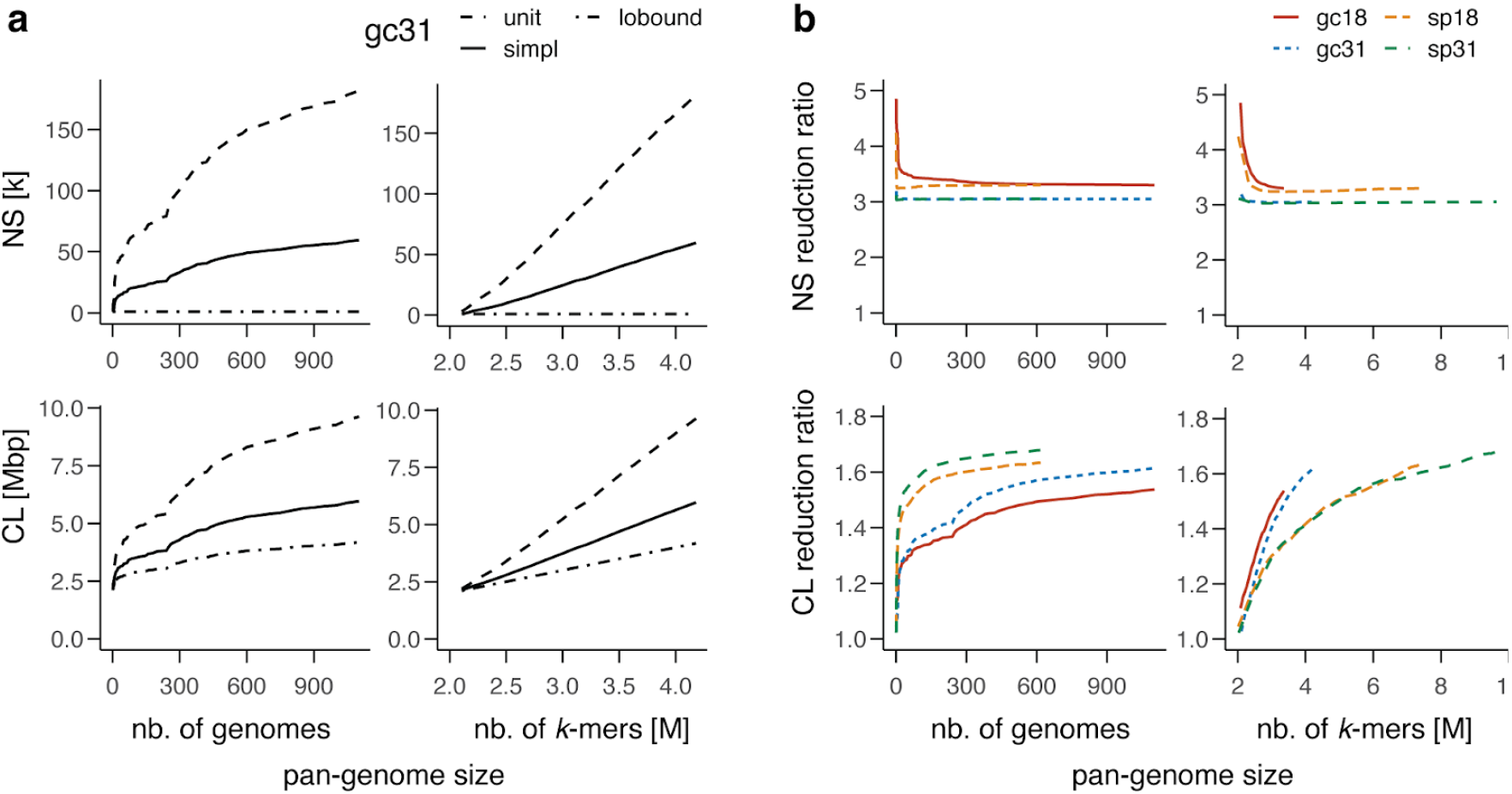
Scaling of simplitigs and unitigs of bacterial pan-genomes as the pan-genome size grows with a better sampling and more within-species variation. **a** Number of sequences (NS, thousands) and their cumulative length (CL, megabase pairs) for simplitigs (simpl) and unitigs (unit) and the lower bound (lobound) of *N. gonorrhoeae*and *k* = 31, as a function of the number of genomes (left) and *k*mers (right, millions) included. **b** Reduction ratio of simplitigs over unitigs for *S. pneumoniae*(sp) and *N. gonorrhoeae*(gc) and *k* = 18, 31, as a function of the number of genomes (left) and *k*mers (right, millions) included. Full results are available in **Additional File 3**.

We first constructed a pan-genome of *N. gonorrhoeae*, and characterized unitigs and simplitigs as a function of pan-genome size (**Fig. 4a**). We used 1,102 draft assemblies of clinical isolates from the Prevention’s Gonococcal Isolate Surveillance Project [41], from which we built a series of de Bruijn graphs using an increasing number of genomes. Consistent with previous experiments (**Fig 2ab**, *k* = 31), both representations perform comparably well when only one bacterial genome is included (**Fig. 4a**). However, as the number of genomes or *k*-mers grows, the NS and CL grow as well, but with an increasing gap between unitigs and simplitigs; importantly, the latter stay close to the theoretical lower bounds. When the pan-genome size is measured via the number of genomes included, the CL and NS resemble logarithmic functions for both unitigs and simplitigs (**Fig. 4a**, left-hand column). However, when the number of *k*-mers included is used instead, the NS and CL functions act as affine functions (**Fig. 4a**, right-hand column). This suggests that a pan-genome *k*-mer count and a species-specific slope may be used as the predictors of simplitig performance in future applications.

To analyze the relative benefit of simplitigs with growing de Bruijn graphs, we evaluated the NS and CL reduction ratio of simplitigs over unitigs in different configurations (**Fig. 4b**). We used the same *N. gonorrhoeae*dataset and considered also another dataset of *S. pneumoniae*, consisting of 616 draft Illumina assemblies of isolates from a carriage study of children in Massachusetts, USA [42,43]. For both species and for *k* = 18, 31, we constructed a series of de Bruijn graphs as previously, but this time we visualized the NS and CL reduction ratios. In all cases, the NS reduction ratios eventually stabilized at values close to 3, following an L-shape (*k* = 18) or being almost constant (*k* = 31). The CL reduction ratio admitted approximately a logarithmic dependence on the number of genomes and still resembled a linear dependence on the number of *k*-mers. Overall, these experiments provided further evidence that the benefit of simplitigs over unitigs grows with the increased proportion of branching nodes in a de Bruijn graph or with increasing data in case of pan-genome reference structures.

### Application of simplitigs for *k*-mer search

Finally, we sought to demonstrate the benefit of simplitigs in a real application. A major use of de Bruijn graphs consists in *k*-mer matching, which requires the graphs to act as a membership data structure. As both simplitigs and unitigs are text-based representations, *k*-mer queries can be implemented using an arbitrary full-text index [44], notably a Burrows-Wheeler Transform index [45] (sometimes referred to as an FM-index). Here, we used the index of BWA [46], as one of the best-engineered solutions available, to analyze the impact of unitig replacement by simplitigs.

### Single pan-genome

We first evaluated the simplitig improvement on the same *N. gonorrhoeae* pan-genome (**Fig. 5**). We considered four different *k*-mer lengths *k* = 19, 23, 27, 31 and for each of them, we built three pan-genome representations from the original draft genome assemblies: first, we merged the assemblies as the most straightforward approach to collect all *k*-mers; then we computed pan-genome representations by unitigs and simplitigs. As all the three representations carry the same *k*-mer set, a full-text index built upon them should provide the same results, but with a performance reflecting the differences in NS and CL.

**Fig. 5.**
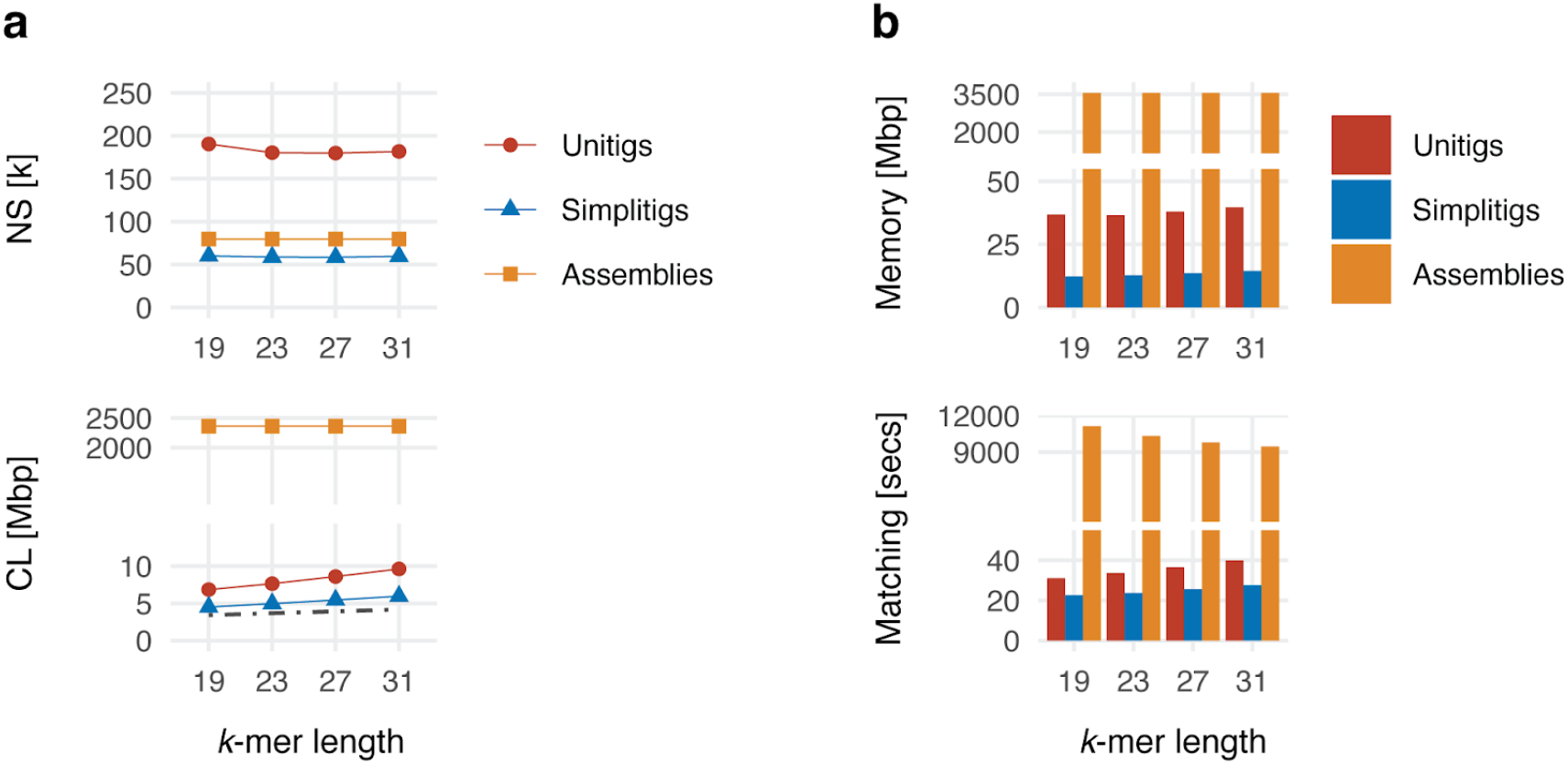
*k*-mer queries for the *N. gonorrhoeae*pan-genome on top of the draft assemblies, unitigs, and simplitigs. **a** Characteristics of the obtained unitigs and simplitigs: number of sequences (NS, thousands) and their cumulative length (CL, megabase pairs). The dot-dash line depicts the CL lower bound corresponding to the number of *k*-mers. **b** Time and memory footprint of BWA *k*-mer queries (10 million *k*-mers). Full results including relative improvements are available in **Additional File 4**.

We analysed the NS and CL characteristics of the computed representations (**Fig. 5a**). Simplitigs improved NS and CL over unitigs by factors of 3.1×–3.2× and 1.5×–1.6×, respectively, consistent with **Fig. 4**. This suggests that memory required for unitigs would be approximately by 50% higher than for unitigs across all the values of *k* considered. We also compared simplitigs and unitigs to assemblies; both improved the CL by two orders of magnitude while the NS stayed comparable for simplitigs and increased twofold for unitigs.

We then evaluated *k*-mer query performance and memory footprint (**Fig. 5b**). For each representation, we constructed a standard BWA index and matched 10 million random *k*-mers from the pan-genome using BWA fastmap [47], with no restriction on maximum size of the suffix-array interval to ensure evaluation correctness (Methods). Surprisingly, simplitigs improved memory by factors of 2.7× – 3.0× (**Fig. 5b**), thus twice as much as we previously anticipated. This is explained by the fact that the underlying full-text engine has to keep information about individual sequences in memory as separate records. As NS grows, it has a negative impact on both the memory footprint and query speed. Nevertheless, since simplitigs provided a substantial reduction in NS over unitigs, this overhead has been reduced. Therefore, the excessive number of unitigs observed throughout our experiments (**Fig. 1** and **Fig. 2**) provides a further argument for replacing unitigs by simplitigs when possible.

### Multiple pan-genomes

We evaluated the performance of the simplitig representation for simultaneous indexing of a large number bacterial pan-genomes (**Fig. 6**). We downloaded all complete bacterial genomes from Genbank that had not been excluded from RefSeq (as of May 2020; 9,869 records out of which 9,032 had genomic sequences available; Methods). We restricted ourselves to complete genomes as draft genomes in Genbank are substantially impacted by false genetic variability [48–50] that is particularly common in bacterial studies, mainly due to the contaminant DNA [51]. By grouping individual genomes per species, we obtained 3,179 bacterial pan-genomes which we call the “All” dataset. After computing simplitigs and unitigs per species, we merged the obtained representations and constructed indexes using BWA; all this was done for *k* = 19, 23, 27, 31 to evaluate the impact of the *k*-mer length. As the unitig index could not fit into RAM of our desktop computer for any *k*, we also created the “Solid” dataset by omitting pan-genomes with less than 11 genomes; this resulted in 112 pan-genomes with 3,958 genomes. We provide all the constructed pan-genomes in the form of simplitigs on Zenodo (<u>10.5281/zenodo.3800713</u>).

**Fig. 6.**
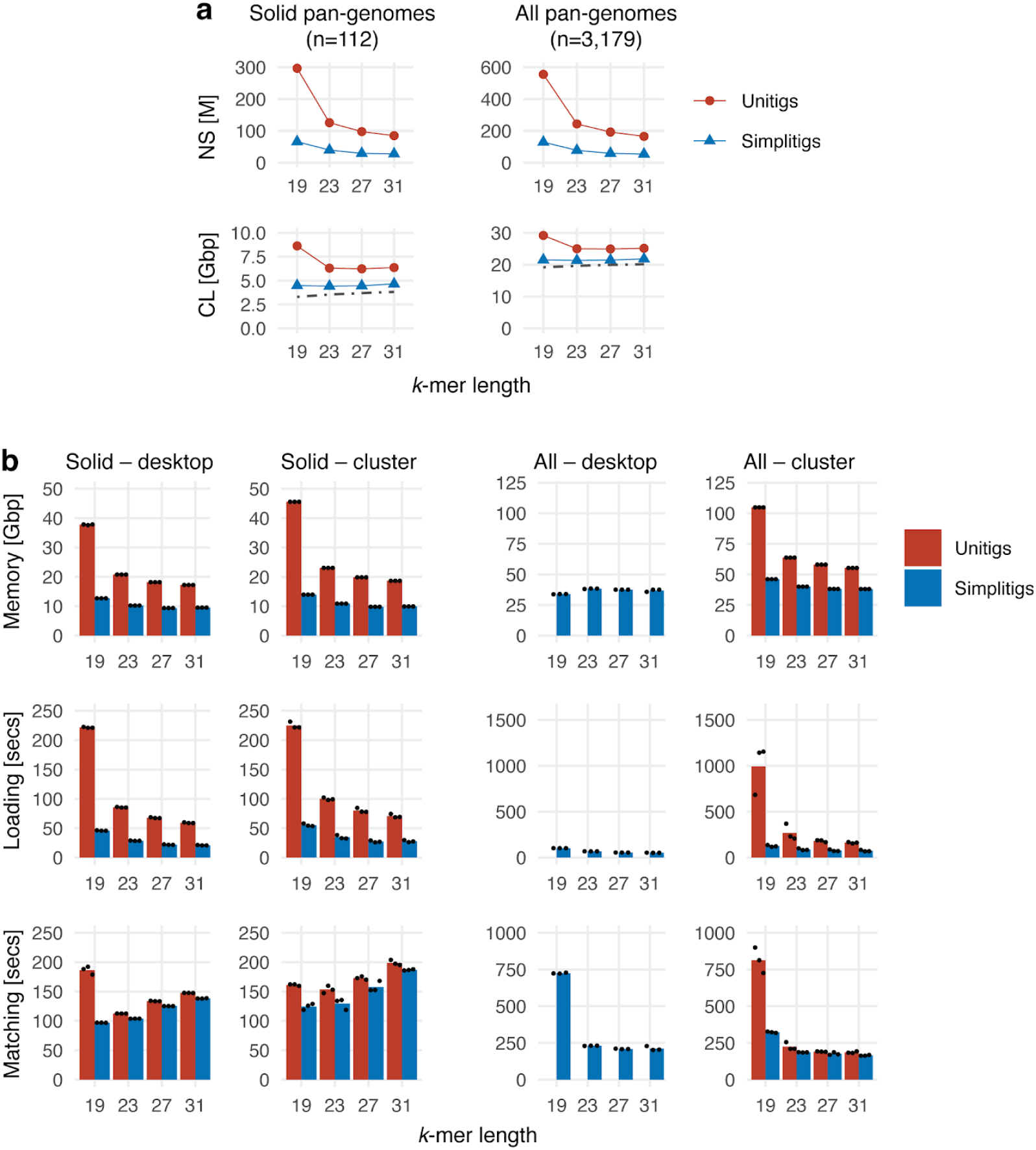
*k*-mer queries for multiple pan-genomes indexed simultaneously. Bacterial pan-genomes were computed from the complete Genbank assemblies per individual species. While the All dataset comprises all pan-genomes with no restriction on their size, the Solid dataset comprises only those that contain at least 11 genomes. **a** Characteristics of the obtained unitigs and simplitigs: number of sequences (NS, millions) and their cumulative length (CL, gigabase pairs). The dot-dash line depicts the lower bound corresponding to the number of *k*-mers. **b** Memory footprint, time of index loading and time of matching 10 million *k*-mers using BWA. The bars correspond to the mean of three measurements (black dots). Full results including relative improvements are available in **Additional File 5**.

First we analyzed the obtained simplitig and unitig representations of both datasets (**Fig. 6a**). We observe that simplitigs provide substantial improvement in both test characteristics. In the Solid dataset, NS and CL were reduced by simplitigs by factors of 3.1–4.5 and 1.4–1.9, respectively; and in the All dataset, NS and CL were reduced by factors of 3.0–4.3 and 1.2–1.4, respectively; all consistent with the scaling observed previously (**Fig. 4**, **Fig. 5**). While the improvement in NS was almost identical in both datasets (consistent with the top-right graph in **Fig 4b**), the improvement in CL was clearly better in the Solid dataset. Indeed, as the vast majority of pan-genomes in the All dataset contained only one genome, the de Bruijn graphs had a comparatively low number of branching nodes, therefore the difference between simplitigs and unitigs was less striking (consistent with the values for small pan-genome sizes in **Fig 4b**). We also observe that, in contrast to unitigs, *k*-mer length had only little impact on the CL of simplitigs within the tested range, which provides better guarantees on required computational resources in future applications.

We then measured the performance of *k*-mer lookup (**Fig. 6b**). Both on a desktop and on a cluster, we evaluated memory footprints, index loading time and time to match ten million random *k*-mers from the index using BWA (Methods). We observed that simplitigs substantially improved the memory footprint and index loading times. For *k* = 19, simplitigs largely improved the matching times, where the difference is caused by “ghost *k*-mers” on unitig borders; these were more common in this experiment due to the short *k*-mer length and the high number of unitigs. For higher *k*-mer lengths, simplitigs still provided a moderate improvement in the matching rate. We note that the query time with BWT-based *k*-mer indexes is dominated by high-frequency *k*-mers. As these are equally frequent in simplitigs and unitigs, the observed performance is similar unless many ghost *k*-mers emerge on sequence borders as seen previously. On the desktop, the unitig All-index could not be evaluated as it did not fit into memory, and the outlier for *k* = 19 may be the result of memory swapping.

## Discussion

We introduced the concept of simplitigs, a generalization of unitigs, and demonstrated that simplitigs constitute a compact, efficient and scalable representation of de Bruijn graphs for various types of genomic datasets. The two representations share many similarities: they are text-based and individual strings correspond to spellings of vertex-disjoint paths. Both representations can be seen as irreversible transforms, taking a set of input strings and producing a new set of strings preserving the *k*-mer sets. In both cases, the resulting files can easily be manipulated using common Unix tools, compressed using standard compression techniques, and indexed using full-text indexes. The main difference consists in that simplitigs do not explicitly carry information about the topology of the de Bruijn graph. Also simplitigs are not expected to have direct biological significance – neighboring segments of the same simplitig may correspond to distant parts of the same nucleic acid or even to different ones. Nevertheless, unitigs can always be recomputed from simplitigs, but this step is not required for many common applications. Furthermore, a concept analogous to simplitigs, called disjointigs, was recently introduced in the context of genome assembly using A-Bruijn graphs [52,53], suggesting that simplitigs may be useful beyond the context of topology-oblivious applications.

We provided ProphAsm, a tool implementing a greedy heuristic to efficiently compute maximal simplitigs from a *k*-mer set. ProphAsm is a spin-off of the ProPhyle software (https://prophyle.github.io, [22,54]) for metagenomic classification, allowing efficient indexing of *k*-mers propagated to individual nodes of a phylogenetic tree. ProphAsm presents a “naive” implementation of the greedy heuristic (**Alg 1)** that can be further improved. For instance, a hash table with better memory management may reduce the memory requirement by a factor of 2.5 [55]. Additional memory reduction may be also achieved in a similar fashion as previously done for unitigs [38,56,57]. Nevertheless, on the studied data, ProphAsm strikingly outcompeted BCALM in all characteristics measured, with the only exception of memory in the case of *H. sapiens*.

The observed lower requirements of ProphAsm can be attributed to two major differences between unitigs and simplitigs. First, unitigs, and BCALM as one of the reference programs, are designed for assembly applications, where only a small proportion of nodes is usually branching. Moreover, experiments are usually run on a cluster in series, with multiple threads per experiment, rather than in parallel, and the tools can use extensive disk space. However, when unitigs are tested as a general representation with restricted disk space, the required resources grow substantially. Second, simplitigs are likely to be, from a computational perspective, fundamentally easier than unitigs.

A challenging but also promising feature of simplitig representation is the ambiguity of maximal simplitigs. This is in sharp contrast to maximal unitigs, which are uniquely defined (up to the order, reverse complementing, and cycles). In practice, every algorithm for simplitig computation has to decide which edge will be included at each branching node. Here, we prioritized speed, simplitigs were constructed progressively and lexicographically minimal edges were used in a case of ambiguity. Therefore, the final maximal simplitigs were only dependent on the choice of seeding *k*-mers; these are determined by the specific implementation of ‘std::unordered_set’ in the C++ standard library. Nevertheless, characteristics other than speed could readily be prioritized instead. For instance, a more sophisticated heuristic may shift the CL and NS closer to the optimum. One could also aim at adding biological meaning to simplitigs, e.g., by preferring those paths that are better supported by sequence data. On the other hand, streaming algorithms for operations such as merging or intersecting may require specific prescribed forms of unitigs. Finally, simplitigs may be optimized for entropy in order to maximize their compressibility.

The data presented in this paper highlight the scaling of computational resources as more sequencing data become available. The studied *N. gonorrhoeae* dataset constitutes a relatively complete image of a bacterial population in a geographical region at a given time scale. As such, it can be used to model the “state of completion” of *k*-mer-based pan-genome representations. On the other hand, the multiple pan-genomes experiment shows how simplitigs perform when a large number of such pan-genomes, although in different states of completion, are considered simultaneously and queried using a single BWT index. Overall, the presented experiments allow us to predict the resources for species for which only limited sequence data are available at present, but more are likely to be generated in the future. Importantly, with more data available, comparative benefits of simplitigs over unitigs grow as we have shown throughout the paper. As the growth of public databases negatively impacts the accuracy of those algorithms that operate on top of lossily represented de Bruijn graphs [58], simplitigs provide a promising solution offering both exactness and scalability.

In modern bioinformatics applications multiple de Bruijn graphs are often considered simultaneously; the resulting structure is usually referred to as a colored de Bruijn graph [14] and the associated data structures have been widely studied [59–70]. Although we touched upon this setting in the Multiple pan-genomes section, exploiting the similarity between individual de Bruijn graphs for further compression in simplitig-based approaches is to be addressed in future work. In many applications, including some of the traditional alignment-free methods [13,71], it is also desirable to consider *k*-mers with counts. In the context of de Bruijn graphs, this leads to the so-called weighted variant of the problem [72]. The fact that frequencies of overlapping *k*-mers are usually similar suggests that *k*-mers can be grouped based on frequencies and simplitigs constructed per group.

Independently and simultaneously with the work we present here, the simplitig representation was recently studied in [73] under the name “spectrum-preserving string sets”. The associated UST tool follows a similar greedy strategy to ProphAsm, although operating on unitigs constructed by BCALM rather than on the original *k*-mers. As we demonstrated throughout this paper, unitigs are prohibitive for highly branching de Bruijn graphs, where a simplitig construction through unitigs may create a burden on resources and easily become intractable. The paper presents a tighter lower bound on the cumulative length of the representation (CL), termed weight, taking into account the graph topology but requiring a computational overhead. The authors also studied the compression properties of simplitigs when combined with standard compression algorithms. On the other hand, the paper does not study the number of sequences (NS) and continuous scaling for parameters such as *k*-mer length or amount of additional variation included.

## Conclusions

In this paper, we addressed the question of efficient and scalable representation of de Bruijn graphs. We showed that the state-of-the-art unitig representation may require adequately large computational resources, especially when de Bruijn graphs contain many branching nodes. We introduced simplitigs, which provide a more compact replacement in applications that do not require explicit information on the graph topology, such as alignment-free sequence comparison and *k*-mer indexing. We introduced a heuristic simplitig computation and showed on the examples of model species that unless the genome is large, even a naive implementation outperforms BCALM, the main state-of-the-art tool for unitigs. We then studied applications to bacterial pan-genomics and showed that the utility of simplitigs compared to unitigs grows as more data are available. Finally, we demonstrated on the example of full-text *k*-mer indexing that simplitigs can substantially reduce computational resources and allow computations in situations where unitigs would bring unaffordable costs when exactness should be preserved.

Our work opens many questions and future directions. The presented algorithm for simplitig computation can be improved, parallelized, de-randomized to ensure reproducibility. We anticipate more theoretical advances in the analysis of the minimum vertex-disjoint path cover problem and better mapping to results from other disciplines such as network sciences. We also anticipate improvements in the heuristic approaches that could simplify and parallelize simplitig computation. The nature of the algorithm suggests that simplitigs might be computed online directly from a stream of data such as sequencing reads. We anticipate better implementations and libraries for simplitigs that can be plugged into standard bioinformatics libraries for various programming languages. Another series of questions is related to low-memory transformations of computed simplitigs that would allow precomputing simplitigs on computer clusters, and tailoring to specific applications on standard computers; this includes decreasing *k*, performing set operations on top of simplitig sets and computing maximal unitigs from simplitigs. A substantial body of work can be anticipated in the direction of text indexing – we showed that simplitigs can be combined with full-text indexes; however, specialized simplitig indexes exploiting simplitig characteristics are yet to be developed. On the theoretical side, many perspectives are open in the direction of orthogonal algorithmic techniques, such as sketching [74,75], and in relation to other stringology concepts, such as minimal absent words [76] and shortest superstrings [77]. Finally, simplitigs can serve as components in design of various specialized data structures; these can involve membership queries of classes of de Bruijn graphs and colored de Bruijn graphs, where simplitigs may encode not only the *k*-mers themselves, but also additional metadata such as *k*-mer frequencies or colors. Furthermore, simplitigs will facilitate new full-text-based data structures for approximate matching, based on the inclusion of the proximity variation, that would be completely intractable with unitigs. Overall, we anticipate that the simplitig representation will become a generic compact representation of de Bruijn graphs, in particular, in the context of large-scale sequence data search engines [69] and sequence data repositories such as those of NCBI and EBI.

## Methods

### De Bruijn graphs

All strings are assumed to be over the alphabet {*A*, *C*, *G*, *T*}. A *k*-mer is a string of length *k*. For a string *s* = *s*_1_…*s_n_*, we define *pref_k_*(*s*) = *s*_1_…*s_k_* and *suf_k_*(*s*) = *s*_*n*−*k*+1_…*s_n_*. For two strings *s* and *t* of length at least *k*, we define the binary connectivity relation *s*→_*k*_*t* if and only if *suf_k_*(*s*) = *pref_k_*(*t*). If *s*→_*k*_*t*, we define the *k*-merging operation ⊙ as *s*⊙^*k*^*t* = *s*· *suf*_|*t*|−*k*_(*t*).

Given a set *K* of *k*-mers, the *de Bruijn graph* of *K*is the directed graph *G*= (*V*, *E*) with *V* = *K* and *E* = {(*u*, *v*) ∈ *K*^2^ | *u*→_*k* − 1_*v*}. For every path *p* = (*v*_1_,…,*v*_*p*_) in *G*, the string *v*_1_⊙^*k*−1^*v*_2_⊙^*k*−1^…⊙^*k*−1^*v_p_* is called a *spelling* of *p*. This definition of de Bruijn graphs is *node-centric*, as nodes are identified with *k*-mers and edges are implicit. Therefore, we can use the terms “ *k*-mer set” and “de Bruijn graph” interchangeably.

### Simplitigs

Consider a set *K* of *k*-mers and the corresponding de Bruijn graph *G*= (*K*, *E*). A *simplitig graph G*′ = (*K*, *E*′) is a spanning subgraph of *G*that is acyclic and the in-degree and out-degree of any node is at most one. It follows from this definition that a simplitig graph is a vertex-disjoint union of paths, whose spellings we call *simplitigs*. A simplitig is called *maximal*if it cannot be extended forward or backward without breaking the definition of simplitig graph. In more detail, a simplitig *u*_1_ →_*k*−1_*u*_2_→_*k*−1_…→_*k*−1_*u_n_*is maximal if the following conditions hold

- either *u*_1_ has no incoming edges in *G*, or for any edge (*v*, *u*_1_) ∈ *E*, *v* belongs to another simplitig and it is not its last vertex,
- either *u_n_* has no outgoing edges in *G*, or for any edge (*u_n_*, *v*) ∈ *E*, *v* belongs to another simplitig and it is not its first vertex.

A *unitig* is a simplitig *u*_1_ →_*k*−1_*u*_2_→_*k*−1_…→_*k*−1_*u_n_* such that each of the nodes *u*_2_,…, *u_n_* has in-degree 1 and each of the nodes *u*_1_,…, *u*_*n*−1_ has out-degree 1 in graph *G*. A maximal unitig is defined similarly.

### Comparing simplitig and unitig representations

Simplitigs and unitigs representations were compared in terms of the number of sequences produced (NS) and their cumulative length (CL). For any set of simplitigs (i.e., not necessarily maximal ones), NS is bounded by 1 and #*kmers*, CL is bounded by #*kmers* and *k*· #*kmers*. The upper bound corresponds to the state of maximal fragmentation, where every *k*-mer forms a simplig. The lower bound corresponds to the maximum possible degree of compaction, i.e., a single simplitig containing all *k*-mers.

NS and CL are readily connected by the following formula:

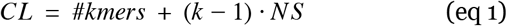

As an important consequence, both characteristics are optimized simultaneously.

### Greedy computation of simplitigs

The problem of computing maximal simplitigs that are optimal in CL (i.e., also in NS) corresponds to the minimum vertex-disjoint path cover problem [78]. This is known to be NP-hard in the general case, reducing from the Hamiltonian path problem. However, the complexity for de Bruijn graphs remains an open question. A greedy heuristic to compute maximal simplitigs has been used throughout this paper (**Alg. 1**). Simplitigs are constructed iteratively, starting from (arbitrary) seeding *k*-mers and being extended greedily forwards and backwards as long as possible.

### ProphAsm implementation

ProphAsm is written in C++ and implements the greedy approach described above (**Alg. 1**). *k*-mers are encoded using uint64_t and stored in an std::unordered_map. The choice of extension nucleotides on branching nodes is done based on the lexicographic order. Therefore, the only source of randomness is the choice of seeding *k*-mers by std::unordered_set::begin; the C++ standard library makes no guarantees on which specific element is considered the first element. ProphAsm does not require any disk space to store intermediate data and its memory consumption corresponded to 38–51 bytes per a unique *k*-mer (in dependence on the allocation), consistent with [55].

### Uni-directed and bi-directed models

The uni-directed model, as presented above, is useful for introducing the concepts of unitigs and simplitigs, but is not directly applicable to data obtained using sequencing: since DNA is double-stranded, every string may come from either strand. At the level of *k*-mers, double-strandedness can be accounted for by using canonical *k*-mers, i.e., by pairing-up every *k*-mer with its reverse complement, typically done by taking the lexicographical minimum of the *k*-mer and its reverse complement. This subsequently requires redefinining de Bruijn graphs to bi-directed de Bruijn graphs [79], which requires a more complex formalism.

### Correctness evaluation

The correctness of simplitig computation can be verified using an arbitrary *k*-mer counter. Simplitigs have been computed correctly if and only if every *k*-mer is present exactly once and the number of distinct *k*-mers is the same as in the original datasets. The correctness of ProphAsm outputs was verified using JellyFish 2 [29].

### Experimental evaluation – model organisms and performance

Reference sequences for six selected model organisms were downloaded from RefSeq and UCSC Genome Browser: *S. pneumoniae* str. ATCC 700669 (accession: NC_011900.1, length 2.22 Mbp), *E. coli* str. K-12 (accession: NC_000913.3, length: 4.64 Mbp), *S. cerevisiae* (accession: NC_001133.9, length: 12.2 Mbp), *C. elegans* (accession: GCF_000002985.6, length: 100 Mbp), *B. mori* (accession: GCF_000151625.1, length: 482 Mbp), and *H. sapiens* (HG38, http://hgdownload.soe.ucsc.edu/goldenPath/hg38/bigZips/hg38.fa.gz, length: 3.21 Gbp). For each genome, simplitigs and unitigs were computed using ProphAsm and BCALM, respectively, for a range of *k*-mer lengths [11,31].

Individual experiments were run in parallel on the Harvard Medical School O2 cluster using Snakemake [80] and SLURM. ProphAsm and BCALM were run with the following parameters, respectively: ‘-k {kmer-length}’ and ‘-kmer-size {kmer-length}-abundance-min 1-nb-cores {cores}-max-disk 30000’. As BCALM requires a large undocumented amount of disk space, we used the -max-disk parameter to make a parallel execution of many BCALM jobs feasible. The SLURM specifications of resource allocation for individual species were iteratively adjusted until all jobs would finish; the final required resources are provided in **Supplementary Table 1**. Time and memory consumption of jobs were measured independently using GNU Time. Individual jobs were deployed to computational nodes with different hardware configurations, which are specified in **Additional File 2**.

**Supplementary Table 1.** SLURM resource allocation for ProphAsm and BCALM2 for the performance evaluation.

| Species | ProphAsm (1 core) |  | BCALM (1 core) |  | BCALM (4 cores) |  |
| --- | --- | --- | --- | --- | --- | --- |
|  | mem<br>[GB] | cpu<br>[hours] | mem<br>[GB] | cpu<br>[hours] | mem<br>[GB] | cpu<br>[hours] |
| *S. pneumoniae* | 10 | 1 | 10 | 1 | 10 | 1 |
| *E. coli* | 10 | 1 | 10 | 1 | 10 | 1 |
| *S. cerevisiae* | 10 | 1 | 10 | 1 | 10 | 1 |
| *C. elegans* | 10 | 1 | 10 | 2 | 10 | 2 |
| *B. mori* | 20 | 2 | 30 | 10 | 30 | 2 |
| *H. sapiens* | 120 | 4 | 100 | 48 | 100 | 18 |

### Experimental evaluation – bacterial pan-genomes

First, 1,102 draft assemblies of *N. gonorrhoeae* clinical isolates (collected from 2000 to 2013 by the Centers for Disease Control and Prevention’s Gonococcal Isolate Surveillance Project [41], and sequenced using Illumina HiSeq) were downloaded from Zenodo [81]. Second, 616 draft assemblies of *S. pneumoniae* isolates (collected from 2001 to 2007 for a carriage study of children in Massachusetts, USA [42,43], and sequenced using Illumina HiSeq) were downloaded from the SRA FTP server using the accession codes provided in Table 1 in [43]. For each of these datasets, an increasing number of genomes was being taken and merged, and simplitigs and unitigs computed using ProphAsm and BCALM, respectively. This experiment was performed for *k* = 18 and *k* = 31. To avoid excessive resource usage the functions were evaluated at selected points in an increasing distance: for intervals [10, 100] and [100,+∞] only multiples of 5 and 20 were evaluated, respectively.

### Experimental evaluation – full-text *k*-mer queries

In the single pan-genome experiment, the same 1,102 assemblies of *N. gonorrhoeae* were merged into a single file. ProphAsm and BCALM were then used to compute simplitigs and unitigs, respectively, from this file for *k* = 19, 23, 27, 31. Each of the three obtained FASTA files (assemblies, simplitigs, and unitigs) was used to construct a BWA index, which was then queried for *k*-mers using ‘bwa fastmap-l {kmer-length}’. We used a modified version of BWA fastmap that reports both the time of index loading and the time of querying (http://github.com/karel-brinda/bwa, commit e1f907c). Query *k*-mers were generated from the same pan-genome using WGsim (version 1.10, with the parameters ‘-h 0 -S 42 -r 0.0 -1 {kmer-length} -N 10000000 -e 0’).

For the multiple pan-genome experiment, a list of available bacterial assemblies was downloaded from ftp://ftp.ncbi.nlm.nih.gov/genomes/genbank/bacteria/assembly_summary.txt (2020/05/05). For all assemblies marked as complete (i.e., the “assembly_level” column equal to “Complete genome”) and present in RefSeq (i.e., an empty value in the column “excluded_from_refseq”), directory urls and species names were extracted (n=9,869). These were then used to download the genomes of the isolates using RSync, restricting to genomic sequences only (i.e., files matching ‘*v?_genomic.fna.gz’, n=9,032). The downloaded assemblies were then merged per species in order to collect *k*-mers of individual pan-genomes and used for computing simplitigs and unitigs using ProphAsm and BCALM, respectively. The obtained simplitig and unitig files were then merged per categories (e.g., simplitigs for k=19) and used to construct a BWA index. The obtained indexes were queried for 1o million *k*-mers using BWA fastmap as previously. The *k*-mers were generated from the original assemblies of randomly selected 100 genomes using DWGsim [82] (version 0.1.11, with the parameters ‘-R 0 -e 0 -r 0 -X 0 -y 0 -H -z 42 -m /dev/null -N 10000000 -1 {k} −2 0’); the randomization was performed using ‘sort -R’.

### Computational setup

The experiments were performed on the HMS O2 research high-performance cluster and on an iMac 4.2 GHz Quad-Core Intel Core i7 with 40 GB RAM. The reproducibility of computation was ensured using BioConda [83].

All benchmarking was performed using ProphAsm 0.1.1 (commit ea28b708) and BCALM 2.2.2 (commit febf79a3). Time and memory footprint were measured using GNU Time.

## Declarations

### Ethics approval and consent to participate

Not applicable.

### Availability of data and materials

#### Additional files

**Additional File 1.** Detailed information for the single genome experiment: NS, CL and #kmers for unitigs and simplitigs as a function of *k*for the 6 species: **a** *S. pneumoniae*, **b** *E. coli*, **c** *S. cerevisiae*, **d** *C. elegans*, **e** *B. mori*, and **f** *H. sapiens*.

**Additional File 2.** Detailed information for the performance comparison. **a** CPU time and memory consumption (both measured by GNU Time and Snakemake) as a function of species, method, number of threads, and *k*-mer length, including the used computational node. **b** Hardware specifications for individual computational nodes.

**Additional File 3.** Detailed information for the pan-genome scaling experiment: **a** *N. gonorrhoeae*, *k* = 18; **b** *N. gonorrhoeae*, *k* = 31; **c** *S. pneumoniae*, *k* = 18; **d** *S. pneumoniae*, *k* = 31.

**Additional File 4.** Detailed information for the single pan-genome *k*-mer indexing experiment. **a** Characteristics of the resulting simplitigs and unitigs for *k* = 19, 23, 27, 31. **b** Memory footprint, index loading time and time to query 10 million *k*-mers using BWA.

**Additional File 5.** Detailed information for the multiple pan-genomes *k*-mer indexing experiment. **a** List of all genomes used for building the pan-genomes (accession code, version, species, filename, number of sequences, genome size [bp]); **b** List of species and the number of genomes included. **c** Characteristics of the resulting simplitigs and unitigs of individual pan-genomes for *k* = 19, 23, 27, 31. **d** Characteristics of the resulting simplitigs and unitigs for the All-dataset and Solid-dataset and *k* = 19, 23, 27, 31. **e** Memory footprint, index loading time and time to query 10 million *k*-mers using BWA (individual repetitions).

#### Data

All data generated or analysed during this study are included in this published article and its supplementary information files. The simplitigs of the Human genome (HG38, for *k* = 10, 11,…, 32) and the obtained Genbank pan-genomes (for *k* = 19, 23, 27, 31) are provided on Zenodo under the accessions 10.5281/zenodo.3770419 and 10.5281/zenodo.3800713, respectively. The code used for the analyses is provided on Github (https://github.com/karel-brinda/simplitigs-supplementary).

### Software

ProphAsm is open source, licensed under the MIT License. The program was developed in C++ and its source code is available from Github (http://github.com/prophyle/prophasm). ProphAsm binaries for Linux and OS X are distributed through BioConda [83] (https://bioconda.github.io/recipes/prophasm/README.html). The source code of the version used in this paper was deposited in Zenodo (<u>10.5281/zenodo.3887035</u>).

### Competing interests

No competing interests.

### Funding

KB and MB were partially supported by the David and Lucile Packard Foundation and NIGMS of the National Institutes of Health under award number R35GM133700. GK was partially funded by RFBR, project 20-07-00652, and joint RFBR and JSPS project 20-51-50007.

### Authors’ contributions

KB, MB, GK designed the study, contributed to interpretation of the results, wrote the manuscript, and approved the final manuscript. KB developed the software and performed the data analysis. KB and GK developed the theory.

## Acknowledgements

The authors thank Jasmijn Baaijens and Roman Cheplyaka for careful reading and valuable comments, and Kamil Salikhov and Simone Pignotti for helpful discussions at the initial stage of this project. Portions of this research were conducted on the O2 high-performance compute cluster, supported by the Research Computing Group at Harvard Medical School.

## Notes

### Competing Interest Statement

The authors have declared no competing interest.

